# A single ancestral *ANP32* locus in ticks creates multiple protein variants that all support the Thogoto virus polymerase

**DOI:** 10.1101/2024.09.16.613268

**Authors:** Zhenyu Zhang, Thomas Nipper, Ishmael D. Aziati, Adrianus C. M. Boon, Andrew Mehle

## Abstract

Viral polymerases rely on cellular cofactors to support efficient transcription of viral genes and replication of the viral genome. The RNA-dependent RNA polymerase of influenza virus, an orthomyxovirus, requires cellular ANP32A or ANP32B proteins for genome replication. However, little is known about whether ANP32 proteins are required by other orthomyxoviruses like the tick-borne thogotoviruses. Recent structural studies coupled with functional assays suggested that the Thogoto virus polymerase uses both ANP32A and ANP32B from brown dog ticks. We clarify here that this tick vector encodes a single *ANP32* locus corresponding to *ANP32A*. This single gene produces multiple protein variants through alternative splicing and start-site selection, all of which enhance Thogoto virus polymerase. Thogoto virus polymerase activity is also enhanced by human and chicken ANP32 proteins. Thus, ANP32A is a deeply conserved pro-viral cofactor and Thogoto virus shows remarkable plasticity utilizing ANP32 homologues separated by almost 1 billion years of evolution.

## Matters Arising

Like all RNA viruses, Orthomyxoviruses like influenza virus use a virally encoded RNA-dependent RNA polymerase to direct replication of the viral genome and transcription of viral mRNAs. While the viral polymerase contains all of the enzymatic activities necessary for these functions, it is absolutely dependent on cellular cofactors to perform these activities during infection^1,2^. Proteins in the ANP32 family were recently identified as essential cofactors for the influenza virus polymerase^3,4^. ANP32 proteins stabilize formation of the polymerase dimer platform that performs genome replication^5–7^. Cells lacking functional ANP32 proteins do not support viral polymerase activity or infection by influenza A viruses^8–10^. Further, species-specific differences in sequence or splicing of *ANP32A* and *ANP32B* alter their ability to be used as cofactors^9,11,12^. For example, humans express three *ANP32* genes (*ANP32A, ANP32B* and *ANP32E*) and encode two additional retrogenes of unclear function (tentatively identified as *ANP32C* and *ANP32D*)^13^. Human ANP32A and ANP32B are the only variants that normally function as cofactors for human influenza viruses, but none of the human ANP32 proteins support avian influenza polymerases^3,9,10^. Rather, avian polymerases are adapted to avian *ANP32A*, which encodes a unique exon duplication creating a protein insertion that is sufficient to confer cofactor activity^3^. Thus, the requirement of influenza virus polymerase to exploit ANP32 family members is a key determinant of viral host range and drives evolutionary adaptations in the virus^12,14^.

While the importance of ANP32 family members has been investigated in detail for influenza viruses, little is known about their use by other orthomyxoviruses like the tick-borne thogotoviruses. Recent studies in *Nature Communications* by Xue et al. provided a comprehensive overview of the structure of the polymerase from the species type Thogoto virus^15^. Complementing their structural studies, they used cell-based polymerase activity assays to test conclusions derived from their models. Assays were performed in human cells lacking ANP32 proteins, establishing a clean genetic background to assess the function of heterologous ANP32 proteins. They report that Thogoto virus polymerase activity is enhanced by expression of “ANP32A” or “ANP32B” derived from the brown dog tick (*Rhipicephalus sanguineus*), a known vector host for thogotoviruses^15^. This was surprising to us, as our genetic analysis suggests that ticks encode only a single *ANP32* gene (Fig 1a). Only one locus was identified in the genome of the brown dog tick, and synteny indicates that this is the locus that became *ANP32A* in humans and other influenza virus hosts like chickens and pigs (Fig 1a). We refer to this as *Rsa ANP32A*. Identical results were found in the genome of the lone star tick (*Amblyomma americanum* [*Aam*]), a distantly related tick species that is the arthropod vector for Bourbon virus, another thogotovirus^16,17^. Analogous syntenic blocks for *ANP32B* and *E* were not found in the tick genome (Fig 1a). Synteny was not conserved between human and avian *ANP32B*, and for human and rodent *ANP32C* (not shown), indicating more recent and independent acquisition of these genes.

**Figure 1.**
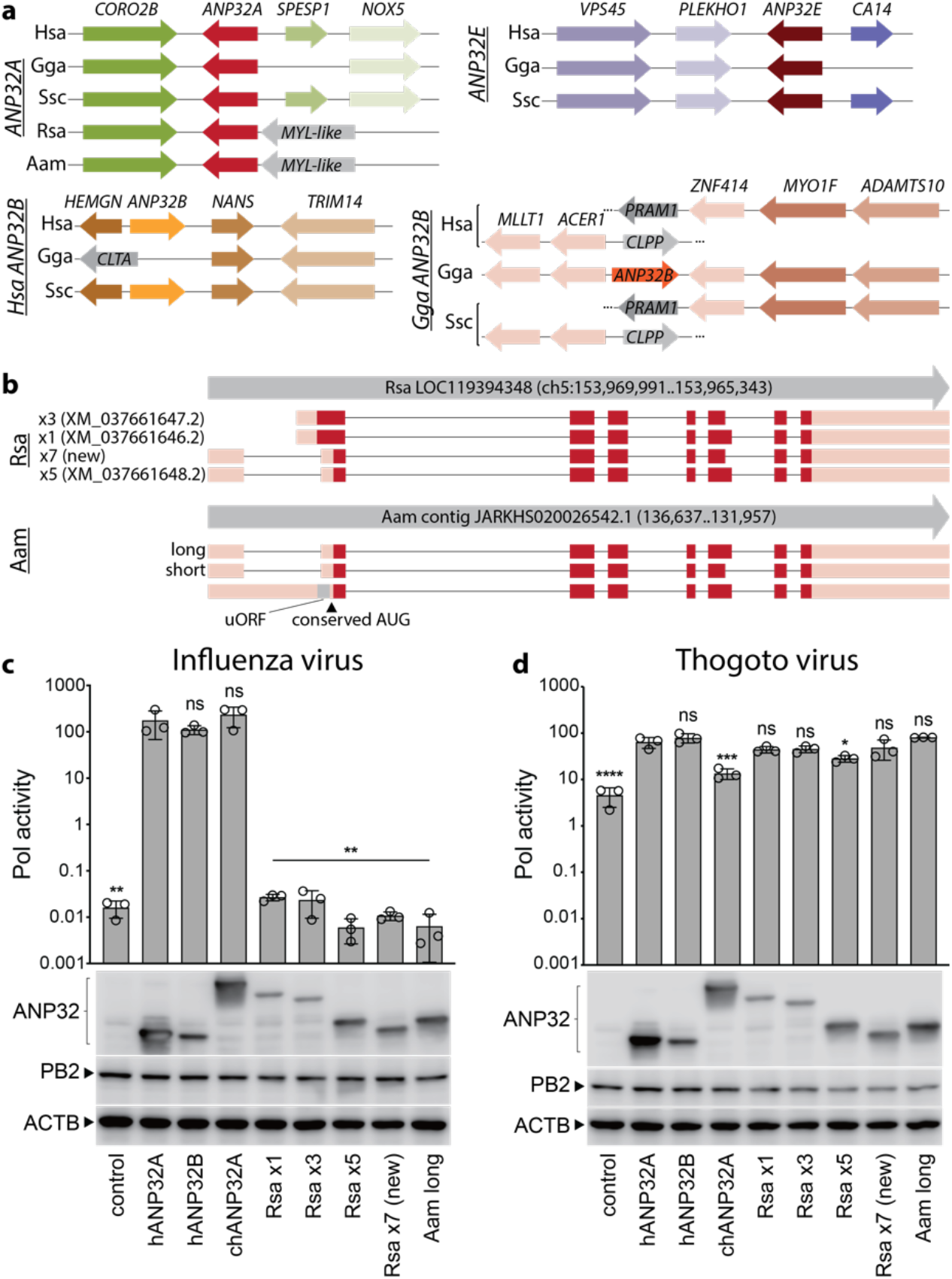
The Thogotovirus polymerase is supported by distinct ANP32 proteins all encoded by the single ancestral locus present in ticks. a) Syntenic gene blocks were identified for ANP32 genes, revealing a single ancestral locus present in the brown dog (Rsa = *Rhipicephalus sanguineus*) and lone star (Aam = *Amblyomma americanum*) ticks. Comparisons were made to humans (Hsa = *Homo sapiens*), chickens (Gga = *Gallus gallus*), and swine (Ssc = *Sus scrofa*). b) *De novo* transcriptome assembly identifies multiple splice variants of tick ANP32. The genomic locus for the brown dog and lone star tick is indicated, as well as accession numbers for previously catalogued transcript variants. Exons are shown as boxes with non-coding regions in pink, open reading frames in red, and a small upstream open reading frame (uORF) in gray. The conserved start site present in vertebrate ANP32A is indicated. c-d) Activity assays demonstrate diverse ANP32 proteins support the Thogotovirus polymerase. Polymerase activity assays were performed in double-knockout human cells lacking ANP32A and ANP32B. c) Influenza virus polymerase, nucleoprotein and a viral reporter were expressed in cells with the indicated human (hANP32), chicken (cANP32) or tick ANP32 protein. Activity was normalized to an internal *Renilla* luciferase control. d) The same approach was used to test activity of the Thogotovirus polymerase. Proteins were detected by western blotting. n = 3 ± sd. Significance was assessed by a one-way ANOVA with a *post hoc* Dunnett’s multiple comparisons test against hANP32A. * = p ≤ 0.05; ** = p ≤ 0.01; *** = p ≤ 0.001; **** = p ≤ 0.0001; ns = not significant.

We looked at the sequence listed under the accession numbers provided by Xue, et al. for brown dog tick “ANP32A” and “ANP32B” that were used in their polymerase activity assays (XP_037517575.1 and XP_037517574.1, respectively). Despite the label suggesting different genes, these are not distinct genes but are instead splice variants that map to the same locus in the tick genome (Fig 1b). We find that automated annotation of the brown dog tick and other tick genomes appears to have resulted in the misidentification of ANP32 proteins – genes that are annotated as distinct *ANP32* family members across many tick species in reality are spliceoforms of ANP32A-like proteins that all map to the same ancestral locus. In Rsa ANP32A, the differential splicing that produced XP_037517575.1 and XP_037517574.1 removed sequences towards the C-terminus of the low complexity acid region that is common to ANP32 proteins. This indel is distinct from the duplication of an exon in chicken ANP32A that confers its species-specific activity^3^. XP_037517575.1 and XP_037517574.1 are also predicted to use an upstream start codon producing a non-canonical extended N-terminus relative to the conserved start codon used in other species. Based on this analysis, the ANP32 proteins used in the functional assays for Thogoto virus polymerase^15^ are two highly unusual variants of the same ANP32 gene.

To identify the actual transcripts and proteins expressed from the ANP32 locus, we performed *de novo* transcriptome assembly using RNA-seq data from the brown dog (Rsa) and lone star (Aam) ticks (Fig 1b, Supplemental Data 1). Data from the brown dog tick transcriptome confirmed that XP_037517575.1 and XP_037517574.1 retain an N-terminal extension and are derived from the x3 and x1 splice variants, respectively. We also identified the previously identified splice variant x5 and a new transcript we classify as x7. x5 and x7 code for canonical ANP32A proteins that have a start codon near the conserved initiation site. Interestingly, Rsa ANP32A proteins that initiate near the conserved start position lack the start codon shared by humans, pigs or chickens, and use a different AUG two codons downstream. Transcripts from the lone star tick also used the downstream start codon near the conserved initiation site and contained splice variants producing distinct low complexity acid regions (Fig 1b). One of the lone star tick transcripts also encoded an AUG 5’ to the conserved start site, but in this case it created a small upstream ORF (uORF) as opposed to a N-terminally extended version of the protein. These results experimentally verified canonical and non-canonical tick ANP32 transcripts and revealed that differential splicing is a deeply conserved feature of the *ANP32A* gene^12,14^.

To test the role of *bona fide* tick ANP32A as a viral cofactor, we performed polymerase activity assays with the Thogoto virus polymerase and the influenza A polymerase as a comparator. We measured polymerase activity in human cells lacking endogenous ANP32A or ANP32B. Cells were complemented with ANP32 proteins from humans, chickens, or ticks. As shown before^9,10^, the influenza virus polymerase is almost completely dependent on the presence of an ANP32 protein, utilizing human and chicken ANP32A and human ANP32B (Fig 1c). However, none of the tick ANP32 variants supported influenza polymerase activity, reflecting the fact that these proteins contain polymorphisms at amino acids 129 and 130 (hANP32A numbering) that are known to prevent their usage as viral cofactors^9,10^. In contrast, the Thogoto virus polymerase retained significant activity in double knockout cells (Fig 1d), mirroring results from Xue et al.^15^ Nonetheless, Thogoto virus polymerase activity was enhanced when supplied with experimentally defined variants from the brown dog tick ANP32A. Canonical versions of ANP32A supported high levels of Thogoto virus polymerase activity (Fig 1d). Brown dog tick ANP32 variants with the unusual extended N-termini also supported the Thogoto virus polymerase, in agreement with the results of Xue, et al. (Fig 1d). ANP32A from the lone star tick also supported high levels of polymerase activity, reinforcing that the N-terminal extension present in brown dog ticks is not essential. Thogoto virus polymerase activity was also enhanced by human ANP32 proteins, and to a lesser degree by chicken ANP32A proteins (Fig 1d). Thus, ANP32A variants in ticks, all expressed from the same gene, serve as host cofactors for the Thogoto virus polymerase.

These data define the repertoire of tick ANP32A proteins that support polymerase activity, revealing that ANP32 is a deeply conserved pro-viral cofactor. They also show that sequence changes, start-site selection, and splice variants all contribute to diversifying proteins expressed from the ancestral ANP32 locus. Finally, they highlight the remarkable plasticity of thogotoviruses, which are able to utilize both human and tick ANP32 proteins that are separated by almost 1 billion years of evolution^18^.

## Methods

Polymerase activity assays were performed following previous protocols^19–21^ by expressing viral polymerase proteins and nucleoprotein with a model genomic RNA encoding firefly luciferase and a plasmid constitutively expressing *Renilla* luciferase as a normalization control. Influenza virus expression vectors were cloned from the A/WSN/1933 strain^22^. Thogoto virus (strain SiAr126) sequences corresponding to PB2 (MT628440.1), PB1 (MT628441.1), PA (MT628442.1) and NP (MT628444.1) genes were synthesized (Twist Biosciences) and cloned into the expression vector pCDNA3.1 using NEBuilder HiFi DNA Assembly Kit (New England Biolabs). Constructs were verified by sequencing. Proteins were detected by western blotting using antibodies recognizing V5 (Proteintech 14440-1-AP) or FLAG (Sigma F3165) epitope tags.

Syntenic *ANP32* gene clusters for humans, chickens, and swine were initially identified in the KEGG Sequence Similarity DataBase (https://www.kegg.jp/kegg/genes.html). These were then used to search for *ANP32* genes and analogous clusters in the reference genome of *Rhipicephalus sanguineus* (BIME_Rsan_1.4). A similar approach was used for the *Amblyomma americanum* genome (ASM3014330v2), although this genome is not fully assembled or annotated yet, thus genes were identified manually in their corresponding contigs.

Tick ANP32A transcripts were identified by using Trinity (v2.15.1) to create *de novo* transcriptomes using RNA-seq data from *Rhipicephalus sanguineus* cell lines (BioProjects PRJNA238793-PRJNA238796)^23,24^. Transcripts were then mapped back to the locus to characterize splice variants. The same approach was used for *Amblyomma americanum* cell lines (BioProjects PRJNA238773,PRJNA238774, PRJNA238776, PRJNA238782, PRJNA238781, PRJNA238784). Open reading frames identified in these transcripts were codon-optimized for expression in humans, synthesized (IDT), cloned into expression vectors with C-terminal epitope tags, and verified by sequencing. Sequences for Rsa and Aam ANP32A variants can be found in Supplemental Data 1.

## Acknowledgments

This work was supported by an H.I. Romnes Faculty Fellow funded by the Wisconsin Alumni Research Foundation to AM, a Vilas Faculty Mid-Career Investigator Award to AM, and the National Institutes of Health/NIAID R01AI164690 (AM), R01AI173327 (ACMB), and T32AI007414 (TN). We thanks Marta Gaglia for critical reading and feedback.

