## Supplemental Data for "A single ancestral *ANP32* locus in ticks creates multiple protein variants that all support the Thogoto virus polymerase"

### Supplemental Data 1

> brown dog Rsa ANP32A x1 (XP_037517574.1)
MYFEGGPHIEVAALLGACFSALPVFEQVGCVPSASAAVVQMEKRIELEKRGKNPEQIHELNLDNCRSTAI
VGLTEEFVNLETLSLINVGLTSLKGFPKLPNLKKLELSDNRISGGLNLLHGSPKLTHLNLSGNKIKGLET
LDPLKEFKNLKNLDLFNCEVTSIENYRDRVFELIPSLKYLDGYDRDEKEAEDSEADDEDGNEEDDEENEV
DGEGESDEDDDDDVNVDDEDDEDGEEDDDVDEEDGEDGEEEVGLDYLEKDDIDEESEGDFYNPYDVDDDE
DGDENESPKGQKRKREEEEDGED

> brown dog Rsa ANP32A x3 (XP_037517575.1)
MYFEGGPHIEVAALLGACFSALPVFEQVGCVPSASAAVVQMEKRIELEKRGKNPEQIHELNLDNCRSTAI
VGLTEEFVNLETLSLINVGLTSLKGFPKLPNLKKLELSDNRISGGLNLLHGSPKLTHLNLSGNKIKGLET
LDPLKEFKNLKNLDLFNCEVTSIENYRDRVFELIPSLKYLDGYDRDEKEAEDSEADDEDGNEEDDEENEV
DGEGESDEDDDDDVNVDDEDDEDGEEDDDVDEEDGEDGEEEEESEGDFYNPYDVDDDEDGDENESPKGQK
RKREEEEDGED

> brown dog Rsa ANP32A x5 (XP_037517576.1)
MEKRIELEKRGKNPEQIHELNLDNCRSTAIVGLTEEFVNLETLSLINVGLTSLKGFPKLPNLKKLELSDN
RISGGLNLLHGSPKLTHLNLSGNKIKGLETLDPLKEFKNLKNLDLFNCEVTSIENYRDRVFELIPSLKYL
DGYDRDEKEAEDSEADDEDGNEEDDEENEVDGEGESDEDDDDDVNVDDEDDEDGEEDDDVDEEDGEDGEE
EVGLDYLEKDDIDEESEGDFYNPYDVDDDEDGDENESPKGQKRKREEEEDGED

> brown dog Rsa ANP32A x7
MEKRIELEKRGKNPEQIHELNLDNCRSTAIVGLTEEFVNLETLSLINVGLTSLKGFPKLPNLKKLELSDN
RISGGLNLLHGSPKLTHLNLSGNKIKGLETLDPLKEFKNLKNLDLFNCEVTSIENYRDRVFELIPSLKYL
DGYDRDEKEAEDSEADDEDGNEEDDEENEVDGEGESDEDDDDDVNVDDEDDEDGEEDDDVDEEDGEDGEE
EEESEGDFYNPYDVDDDEDGDENESPKGQKRKREEEEDGED

>lone star Aam ANP32A long
MEKRIELEKRGKNPEQIHELNLDNCRSTAIVGLTEEFVNLETLSLINVGLTSLKGFPKLPNLKKLELSDN
RISGGLNLLHGSPKLTHLNLSGNKIKGLETLDPLKEFKNLKNLDLFNCEVTSIENYRDRVFELIPSLKYL
DGYDRDEKEAEDSEVDDEDGNEEDDEENEVDGEGESDEDDEDDVNVDDEDDEDGEEDDDVDEEDGEDGEE
EVGLDYLEKDDIDEESEGDFYNPYDVDDDEDGDENESPKGQKRKREEEDGED

>lone star Aam ANP32A short
MEKRIELEKRGKNPEQIHELNLDNCRSTAIVGLTEEFVNLETLSLINVGLTSLKGFPKLPNLKKLELSDN
RISGGLNLLHGSPKLTHLNLSGNKIKGLETLDPLKEFKNLKNLDLFNCEVTSIENYRDRVFELIPSLKYL
DGYDRDEKEAEDSEVDDEDGNEEDDEENEVDGEGESDEDDEDDVNVDDEDDEDGEEDDDVDEEDGEDGEE
EEESEGDFYNPYDVDDDEDGDENESPKGQKRKREEEDGED

>brown dog Rsa ANP32A assembled mRNA x1
GGAGGCCCGAAGAGCCACGTGACGTAGTCCTTCCCCGATCGTTCGCCGCCTGCGATTGGACGTTTTGAAC
GTGACCTCATGAATAGTAATTAGTTTGCTACGGCGTACATGTACTTTGAAGGAGGACCCCATATTGAGGT
TGCCGCGCTGCTTGGCGCCTGCTTTTCAGCGCTGCCTGTGTTCGAGCAAGTCGGCTGCGTGCCATCCGCC
TCTGCTGCTGTCGTGCAAATGGAGAAGAGGATTGAATTGGAGAAGCGGGGTAAAAACCCCGAGCAGATAC
ACGAGCTCAACCTCGACAACTGCAGAAGTACCGCCATCGTCGGCCTCACAGAGGAATTCGTCAACCTCGA
GACGCTCAGCCTCATCAATGTCGGCCTCACGAGTCTCAAGGGCTTCCCCAAGCTTCCCAACCTCAAAAAG
TTGGAGTTGAGCGACAACAGGATCTCGGGCGGCCTCAACCTTCTGCACGGGAGCCCCAAGCTTACGCATT
TAAATTTGAGCGGGAACAAGATCAAAGGGCTCGAGACCTTGGATCCCTTGAAAGAGTTCAAGAACCTAAA
GAACCTCGATCTTTTCAACTGCGAAGTGACGAGCATCGAGAACTACAGGGACCGGGTGTTTGAACTCATC
CCAAGCCTCAAGTACTTGGACGGCTACGACAGAGACGAGAAGGAAGCTGAGGACTCTGAGGCAGACGACG
AGGACGGTAACGAAGAGGACGATGAGGAGAATGAAGTGGACGGAGAGGGAGAGAGTGATGAGGATGACGA

CGACGATGTGAACGTGGACGACGAAGATGATGAGGATGGCGAGGAGGATGACGATGTTGATGAGGAAGAT
GGAGAAGATGGGGAGGAAGAGGTCGGTCTCGATTACCTAGAAAAAGACGACATTGATGAGGAAAGTGAAG
GAGACTTCTACAACCCATACGACGTAGATGACGACGAAGACGGCGACGAGAACGAGTCTCCAAAGGGCCA
GAAGCGAAAGAGGGAAGAGGAGGAGGACGGGGAGGACTGAGGGAGGGCGGGAGAAGCACTCAACACCAAT
TGGAACATCCCGGCATCTTTACTTTAGTTTTTTTTTTAAACGTCTTCGTCATTGTCACCTGTAGACTTTG
TACAGCGGCGGACAGAAACAAAATGCAACATCACAATCGCATCATCCTCGGTACCTGTCCCTAGCTTTAT
TTTCCCCCATTCATTTCATTCTTGAACAAGATTTTTTTTAAGCTTTTGCTTGCTTTGGGTGGCCTCCATT
GGTTACCCACTAGTTTGCATTATTATATAAGCTTTGTATAGGTTGTTACTGGTGAACCAGGTTTTGTCTT
TGGCAACGACTGTGAAAACAAATGGTCACCCTTGCAGCCATGCACATTCCAAGAGCTGTGTGCTGTTTGG
GGGCGCTGCCGTCTTTCTGCATCTGCTTGCCTGTGTGAGGCAGCAAATGCACTTTTGATAACGCAAGGCA
ACACCAAAGTTGGCCCAGAAAAGGGGGTGGCCTCTAGGGACGGTGCCGTCGGGCACATCCTCCTCATCAT
CATCCACACAAATTTTTTTTTTTTTTTTTCTCGATCCGTATTCTTCTTGGCCCTCCCTTTTCTCAATTTG
CTTTCAAACATTGAATGAAATGCCTTTTATTGTCGCATGTTGGCTGCGAGGGAAGTGTCTGTGCAGGTGC
ATATTCTATTGCATGAAAGTTTTGCAGCCGTTCAGAGACTTCCACTGCCGCCGTCAGTCTTCCGCGGCGC
AGAGGCGCCGTGGAAGTGTAAATTATCAGCAAGGGCTTTGTCCAGTGTTTGGTTCAATTGTTAAATAAAG
TTTTTACTGAAATGGCAAGCACCGCTGTTTGTAACCTCACTGAATTAAGAGCGTACCTTCTGTCCATAGT
GCCTTGGCAGATGCAGTTGCTACAGGAACCTCTCCGTGGATTGATGGTAGGGCTATTCCAATGCGCAGCA
AGACGCTAAACACTGGTTAAATGTGTAAACATCGGACTGGTACCACATTGAAGTTGTGGTCGACCACCAA
TTTAGCGTATCTGAAACTGATTCAAAGGTTAGTCTTTCTGCACCCTGTAGTCTGAATTGAGATCCCTGCT
GCTGTTTTGTTCGTGGCTCCAAATTGTTTGGTTCCTCCAATATTGAACATCTAGAGTGGAGACTTCATCT
GC

>brown dog Rsa ANP32A assembled mRNA x3
GGAGGCCCGAAGAGCCACGTGACGTAGTCCTTCCCCGATCGTTCGCCGCCTGCGATTGGACGTTTTGAAC
GTGACCTCATGAATAGTAATTAGTTTGCTACGGCGTACATGTACTTTGAAGGAGGACCCCATATTGAGGT
TGCCGCGCTGCTTGGCGCCTGCTTTTCAGCGCTGCCTGTGTTCGAGCAAGTCGGCTGCGTGCCATCCGCC
TCTGCTGCTGTCGTGCAAATGGAGAAGAGGATTGAATTGGAGAAGCGGGGTAAAAACCCCGAGCAGATAC
ACGAGCTCAACCTCGACAACTGCAGAAGTACCGCCATCGTCGGCCTCACAGAGGAATTCGTCAACCTCGA
GACGCTCAGCCTCATCAATGTCGGCCTCACGAGTCTCAAGGGCTTCCCCAAGCTTCCCAACCTCAAAAAG
TTGGAGTTGAGCGACAACAGGATCTCGGGCGGCCTCAACCTTCTGCACGGGAGCCCCAAGCTTACGCATT
TAAATTTGAGCGGGAACAAGATCAAAGGGCTCGAGACCTTGGATCCCTTGAAAGAGTTCAAGAACCTAAA
GAACCTCGATCTTTTCAACTGCGAAGTGACGAGCATCGAGAACTACAGGGACCGGGTGTTTGAACTCATC
CCAAGCCTCAAGTACTTGGACGGCTACGACAGAGACGAGAAGGAAGCTGAGGACTCTGAGGCAGACGACG
AGGACGGTAACGAAGAGGACGATGAGGAGAATGAAGTGGACGGAGAGGGAGAGAGTGATGAGGATGACGA
CGACGATGTGAACGTGGACGACGAAGATGATGAGGATGGCGAGGAGGATGACGATGTTGATGAGGAAGAT
GGAGAAGATGGGGAGGAAGAGGAGGAAAGTGAAGGAGACTTCTACAACCCATACGACGTAGATGACGACG
AAGACGGCGACGAGAACGAGTCTCCAAAGGGCCAGAAGCGAAAGAGGGAAGAGGAGGAGGACGGGGAGGA
CTGAGGGAGGGCGGGAGAAGCACTCAACACCAATTGGAACATCCCGGCATCTTTACTTTAGTTTTTTTTT
TAAACGTCTTCGTCATTGTCACCTGTAGACTTTGTACAGCGGCGGACAGAAACAAAATGCAACATCACAA
TCGCATCATCCTCGGTACCTGTCCCTAGCTTTATTTCCCCCATTCATTTCATTCTTGAACAAGATTTTTT
TTTAAGCTTTTGCTTGCTTTGGGTGGCCTCCATTGGTTACCCACTAGTTTGCATTATTATATAAGCTTTG
TATAGGTTGTTACTGGTGAACCAGGTTTTGTCTTTGGCAACGACTGTGAAAACAAATGGTCACCCTTGCA
GCCATGCACATTCCAAGAGCTGTGTGCTGTTTGGGGGCGCTGCCGTCTTTCTGCATCTGCTTGCCTGTGT
GAGGCAGCAAATGCACTTTTGATAACGCAAGGCAACACCAAAGTTGGCCCAGAAAAGGGGGTGGCCTCTA
GGGACGGTGCCGTCGGGCACATCCTCCTCATCATCATCCACACAAATTTTTTTTTTTTTTTTTCTCGATC
CGTATTCTTCTTGGCCCTCCCTTTTCTCAATTTGCTTTCAAACATTGAATGAAATGCCTTTTATTGTCGC
ATGTTGGCTGCGAGGGAAGTGTCTGTGCAGGTGCATATTCTATTGCATGAAAGTTTTGCAGCCGTTCAGA
GACTTCCACTGCCGCCGTCAGTCTTCCGCGGCGCAGAGGCGCCGTGGAAGTGTAAATTATCAGCAAGGGC
TTTGTCCAGTGTTTGGTTCAATTGTTAAATAAAGTTTTTACTGAAATGGCAAGCACCGCTGTTTGTAACC
TCACTGAATTAAGAGCGTACCTTCTGTCCATAGTGCCTTGGCAGATGCAGTTGCTACAGGAACCTCTCCG
TGGATTGATGGTAGGGCTATTCCAATGCGCAGCAAGACGCTAAACACTGGTTAAATGTGTAAACATCGGA
CTGGTACCACATTGAAGTTGTGGTCGACCACCAATTTAGCGTATCTGAAACTGATTCAAAGGTTAGTCTT

TCTGCACCCTGTAGTCTGAATTGAGATCCCTGCTGCTGTTTTGTTCGTGGCTCCAAATTGTTTGGTTCCT
CCAATATTGAACATCTAGAGTGGAGACTTCATCTGC

>brown dog Rsa ANP32A assembled mRNA x5
CGCAATGCCTGTTCTGCGCCAGAGCGTTGGCACAGCTGGTGCGGCACGGAGTCGGGCATGTCTCAAAGCC
CTTGCAGTACTGTGGCCGGCATGGCTTGCGCGGACTTGACACTGAATGCGACATTGTCGAAGCCAAGTGC
GGTCAGAAACGTGTCGGGGTGCCACGAAACGAAGACGAAAATTAGTTAGTGCGCGGCAGCCGTGGAAAGC
CTAACAAACAAACATCAGCGATGGCGCCGTTGCGGACCCGTTTAATCGAGAGGTCTCTGTGTAAAGCAAG
CGCTGATATCAACTAGAGAACATGCGAAAGGCACCGCGCTGCCTGTGTTCGAGCAAGTCGGCTGCGTGCC
ATCCGCCTCTGCTGCTGTCGTGCAAATGGAGAAGAGGATTGAATTGGAGAAGCGGGGTAAAAACCCCGAG
CAGATACACGAGCTCAACCTCGACAACTGCAGAAGTACCGCCATCGTCGGCCTCACAGAGGAATTCGTCA
ACCTCGAGACGCTCAGCCTCATCAATGTCGGCCTCACGAGTCTCAAGGGCTTCCCCAAGCTTCCCAACCT
CAAAAAGTTGGAGTTGAGCGACAACAGGATCTCGGGCGGCCTCAACCTTCTGCACGGGAGCCCCAAGCTT
ACGCATTTAAATTTGAGCGGGAACAAGATCAAAGGGCTCGAGACCTTGGATCCCTTGAAAGAGTTCAAGA
ACCTAAAGAACCTCGATCTTTTCAACTGCGAAGTGACGAGCATCGAGAACTACAGGGACCGGGTGTTTGA
ACTCATCCCAAGCCTCAAGTACTTGGACGGCTACGACAGAGACGAGAAGGAAGCTGAGGACTCTGAGGCA
GACGACGAGGACGGTAACGAAGAGGACGATGAGGAGAATGAAGTGGACGGAGAGGGAGAGAGTGATGAGG
ATGACGACGACGATGTGAACGTGGACGACGAAGATGATGAGGATGGCGAGGAGGATGACGATGTTGATGA
GGAAGATGGAGAAGATGGGGAGGAAGAGGTCGGTCTCGATTACCTAGAAAAAGACGACATTGATGAGGAA
AGTGAAGGAGACTTCTACAACCCATACGACGTAGATGACGACGAAGACGGCGACGAGAACGAGTCTCCAA
AGGGCCAGAAGCGAAAGAGGGAAGAGGAGGAGGACGGGGAGGACTGAGGGAGGGCGGGAGAAGCACTCAA
CACCAATTGGAACATCCCGGCATCTTTACTTTAGTTTTTTTTTTAAACGTCTTCGTCATTGTCACCTGTA
GACTTTGTACAGCGGCGGACAGAAACAAAATGCAACATCACAATCGCATCATCCTCGGTACCTGTCCCTA
GCTTTATTTTCCCCCATTCATTTCATTCTTGAACAAGATTTTTTTTAAGCTTTTGCTTGCTTTGGGTGGC
CTCCATTGGTTACCCACTAGTTTGCATTATTATATAAGCTTTGTATAGGTTGTTACTGGTGAACCAGGTT
TTGTCTTTGGCAACGACTGTGAAAACAAATGGTCACCCTTGCAGCCATGCACATTCCAAGAGCTGTGTGC
TGTTTGGGGGCGCTGCCGTCTTTCTGCATCTGCTTGCCTGTGTGAGGCAGCAAATGCACTTTTGATAACG
CAAGGCAACACCAAAGTTGGCCCAGAAAAGGGGGTGGCCTCTAGGGACGGTGCCGTCGGGCACATCCTCC
TCATCATCATCCACACAAATTTTTTTTTTTTTTTTTCTCGATCCGTATTCTTCTTGGCCCTCCCTTTTCT
CAATTTGCTTTCAAACATTGAATGAAATGCCTTTTATTGTCGCATGTTGGCTGCGAGGGAAGTGTCTGTG
CAGGTGCATATTCTATTGCATGAAAGTTTTGCAGCCGTTCAGAGACTTCCACTGCCGCCGTCAGTCTTCC
GCGGCGCAGAGGCGCCGTGGAAGTGTAAATTATCAGCAAGGGCTTTGTCCAGTGTTTGGTTCAATTGTTA
AATAAAGTTTTTACTGAAATGGCAAGCACCGCTGTTTGTAACCTCACTGAATTAAGAGCGTACCTTCTGT
CCATAGTGCCTTGGCAGATGCAGTTGCTACAGGAACCTCTCCGTGGATTGATGGTAGGGCTATTCCAATG
CGCAGCAAGACGCTAAACACTGGTTAAATGTGTAAACATCGGACTGGTACCACATTGAAGTTGTGGTCGA
CCACCAATTTAGCGTATCTGAAACTGATTCAAAGGTTAGTCTTTCTGCACCCTGTAGTCTGAATTGAGAT
CCCTGCTGCTGTTTTGTTCGTGGCTCCAAATTGTTTGGTTCCTCCAATATTGAACATCTAGAGTGGAGAC
TTCATCTGC

>brown dog Rsa ANP32A assembled mRNA x7
CGCAATGCCTGTTCTGCGCCAGAGCGTTGGCACAGCTGGTGCGGCACGGAGTCGGGCATGTCTCAAAGCC
CTTGCAGTACTGTGGCCGGCATGGCTTGCGCGGACTTGACACTGAATGCGACATTGTCGAAGCCAAGTGC
GGTCAGAAACGTGTCGGGGTGCCACGAAACGAAGACGAAAATTAGTTAGTGCGCGGCAGCCGTGGAAAGC
CTAACAAACAAACATCAGCGATGGCGCCGTTGCGGACCCGTTTAATCGAGAGGTCTCTGTGTAAAGCAAG
CGCTGATATCAACTAGAGAACATGCGAAAGGCACCGCGCTGCCTGTGTTCGAGCAAGTCGGCTGCGTGCC
ATCCGCCTCTGCTGCTGTCGTGCAAATGGAGAAGAGGATTGAATTGGAGAAGCGGGGTAAAAACCCCGAG
CAGATACACGAGCTCAACCTCGACAACTGCAGAAGTACCGCCATCGTCGGCCTCACAGAGGAATTCGTCA
ACCTCGAGACGCTCAGCCTCATCAATGTCGGCCTCACGAGTCTCAAGGGCTTCCCCAAGCTTCCCAACCT
CAAAAAGTTGGAGTTGAGCGACAACAGGATCTCGGGCGGCCTCAACCTTCTGCACGGGAGCCCCAAGCTT
ACGCATTTAAATTTGAGCGGGAACAAGATCAAAGGGCTCGAGACCTTGGATCCCTTGAAAGAGTTCAAGA
ACCTAAAGAACCTCGATCTTTTCAACTGCGAAGTGACGAGCATCGAGAACTACAGGGACCGGGTGTTTGA
ACTCATCCCAAGCCTCAAGTACTTGGACGGCTACGACAGAGACGAGAAGGAAGCTGAGGACTCTGAGGCA

GACGACGAGGACGGTAACGAAGAGGACGATGAGGAGAATGAAGTGGACGGAGAGGGAGAGAGTGATGAGG
ATGACGACGACGATGTGAACGTGGACGACGAAGATGATGAGGATGGCGAGGAGGATGACGATGTTGATGA
GGAAGATGGAGAAGATGGGGAGGAAGAGGAGGAAAGTGAAGGAGACTTCTACAACCCATACGACGTAGAT
GACGACGAAGACGGCGACGAGAACGAGTCTCCAAAGGGCCAGAAGCGAAAGAGGGAAGAGGAGGAGGACG
GGGAGGACTGAGGGAGGGCGGGAGAAGCACTCAACACCAATTGGAACATCCCGGCATCTTTACTTTAGTT
TTTTTTTTAAACGTCTTCGTCATTGTCACCTGTAGACTTTGTACAGCGGCGGACAGAAACAAAATGCAAC
ATCACAATCGCATCATCCTCGGTACCTGTCCCTAGCTTTATTTTCCCCCATTCATTTCATTCTTGAACAA
GATTTTTTTTAAGCTTTTGCTTGCTTTGGGTGGCCTCCATTGGTTACCCACTAGTTTGCATTATTATATA
AGCTTTGTATAGGTTGTTACTGGTGAACCAGGTTTTGTCTTTGGCAACGACTGTGAAAACAAATGGTCAC
CCTTGCAGCCATGCACATTCCAAGAGCTGTGTGCTGTTTGGGGGCGCTGCCGTCTTTCTGCATCTGCTTG
CCTGTGTGAGGCAGCAAATGCACTTTTGATAACGCAAGGCAACACCAAAGTTGGCCCAGAAAAGGGGGTG
GCCTCTAGGGACGGTGCCGTCGGGCACATCCTCCTCATCATCATCCACACAAATTTTTTTTTTTTTTTT
CTCGATCCGTATTCTTCTTGGCCCTCCCTTTTCTCAATTTGCTTTCAAACATTGAATGAAATGCCTTTTA
TTGTCGCATGTTGGCTGCGAGGGAAGTGTCTGTGCAGGTGCATATTCTATTGCATGAAAGTTTTGCAGCC
GTTCAGAGACTTCCACTGCCGCCGTCAGTCTTCCGCGGCGCAGAGGCGCCGTGGAAGTGTAAATTATCAG
CAAGGGCTTTGTCCAGTGTTTGGTTCAATTGTTAAATAAAGTTTTTACTGAAATGGCAAGCACCGCTGTT
TGTAACCTCACTGAATTAAGAGCGTACCTTCTGTCCATAGTGCCTTGGCAGATGCAGTTGCTACAGGAAC
CTCTCCGTGGATTGATGGTAGGGCTATTCCAATGCGCAGCAAGACGCTAAACACTGGTTAAATGTGTAAA
CATCGGACTGGTACCACATTGAAGTTGTGGTCGACCACCAATTTAGCGTATCTGAAACTGATTCAAAGGT
TAGTCTTTCTGCACCCTGTAGTCTGAATTGAGATCCCTGCTGCTGTTTTGTTCGTGGCTCCAAATTGTTT
GGTTCCTCCAATATTGAACATCTAGAGTGGAGACTTCATCTGC

>lone star Aam ANP32A assembled mRNA long
AAATATCCCCTCGATGCTATGCCTGTTTTGCGGCAGAGCGTTTTCAGTAAGCGCGACATAGAGTTCGGCG
TGTCCTACGGTTCCGTTGCAACGGTGGCTGGCATGGCTCGCGCGGACTCCACACTGAATGCGACATTTTC
GAAGCAAGGTGTGCTCGGAAACGTGCCGAGGTGCCACGAAACGAAGACGAAAATTAGTTAGTGCTCGGCG
AACGAGGCAAGCCTGTAGTAACAAACAAACAGCAGCGATGGCGGCGTTTGGCGTTGCGGACTGTTTAGTC
GAGAACTCTCGGTATAAAGCAAGCGCTGATATCAACTAGAGAACATGCGAAAGGCACGGAGCTGCTTGTG
TGTTCTACCGAGTCGGGTGCGTGCCATCCGCCTGTGGTGCTGTAGTGCAAATGGAGAAGAGGATTGAATT
GGAGAAGAGGGGTAAAAACCCCGAGCAGATACACGAGCTCAACCTCGACAACTGCAGGAGTACCGCCATC
GTCGGCCTGACAGAGGAGTTCGTCAACTTGGAGACGCTCAGCCTCATCAATGTCGGCCTTACGAGCCTCA
AGGGCTTCCCCAAGCTGCCCAACCTCAAGAAGTTGGAGCTTAGCGACAACAGGATTTCTGGTGGCCTCAA
CCTGCTGCACGGGAGCCCCAAGCTCACACATTTAAATTTGAGCGGGAACAAAATCAAGGGTCTGGAGACC
TTGGATCCTCTGAAAGAGTTCAAGAACCTTAAGAACCTTGATCTTTTCAACTGTGAAGTGACAAGCATTG
AGAACTACAGAGACAGAGTCTTCGAACTTATCCCGAGCTTGAAGTACCTGGACGGCTATGACCGAGATGA
GAAGGAGGCAGAGGACTCCGAGGTGGACGACGAAGATGGCAATGAGGAGGATGACGAGGAGAATGAAGTG
GATGGGGAGGGAGAGAGTGATGAGGACGATGAAGATGATGTAAACGTGGACGACGAAGATGATGAGGATG
GGGAGGAAGATGACGACGTCGATGAGGAGGATGGAGAAGATGGGGAGGAAGAGGTCGGTCTCGATTACCT
TGAAAAAGACGACATTGATGAGGAAAGTGAAGGAGATTTCTACAACCCATATGATGTGGATGATGACGAA
GATGGAGATGAAAACGAGTCTCCAAAAGGCCAGAAGCGAAAGCGGGAAGAGGAAGATGGGGAGGACTGAG
GGAGGGTGGGAAGAGCACTCAACACAACTGGAACATCCCGGCATCTTTACTTTAGTTTTTTTTTATGTCT
TCTTCATTGTCACCTGTAGACTTTGTACAGCGGCAGAGAGAAACAAAATGCAACATCACAATCGCATCAT
CCTCGGTACCTGTCCCTAGCCTTTCTTCCCCCTTGTTCATTTCATTTGTGAACAATTTTTTGTGTGTGTG
TGTTTTAAAAAGCTTTTGCATGCATGGATTCCCGTTAAGATGCCCACCCAGTTGCTTCGAGATTTGTATA
GGTTGTTACTGCTAAACCAGATTTGATTTCTGGCTACGACCGCAATCAATCGTGCTTACAGCCATGCACA
TTCCAAGAGCTGTGCTGTTACGGGCGCTTTTGTCTTTTGCATCTGCGCCTTCCTCAATCTGAGGGACATC
AAGTTAACTTTGAAAGCAGCGACACCAAAGTTGGCCCAGAAAAGGGGGTGGCCCCTAGGGACAGTGCTGT
GCAGGCACATCTACATCGTCATCCGCGTTTTTTTTTTTCTTTCTCCTTGGCCCTCCCTTTTCTCGATTTG
CTTTCAAAACATTGAATGAAATGCCTTTTATTGTCGCATTGTTGGCTTCGAGGGAAGGGCCTGTGCGGGT
GCACTGCACTTTTAATGTGAGCGACCGTGCGGGCGCTTCCGTTGCGGCTGCCAGTCTTCTGTGGCACAGA
GGTGCGCAGAAGTGTAAATTATCAGCAGGGGCTTTGTCCAGTGTTTGGTTCAATTGTTAAATAAAGTTTT
TACTGAAATGGCGAAAAAAAAAA

>lone star Aam ANP32A assembled mRNA long v2
AAATATCCCCTCGATGCTATGCCTGTTTTGCGGCAGAGCGTTTTCAGTAAGCGCGACATAGAGTTCGGCG
TGTCCTACGGTTCCGTTGCAACGGTGGCTGGCATGGCTCGCGCGGACTCCACACTGAATGCGACATTTTC
GAAGCAAGGTGTGCTCGGAAACGTGCCGAGGTGCCACGAAACGAAGACGAAAATTAGTTAGTGCTCGGCG
AACGAGGCAAGCCTGTAGTAACAAACAAACAGCAGCGATGGCGGCGTTTGGCGTTGCGGACTGTTTAGTC
GAGAACTCTCGGTATAAAGCAAGCGCTGATATCAACTAGAGAACATGCGAAAGGCACGGGTAAGCGTTTT
TCAAGTTATTCGCGCACAGAGCATAGCTAGAAGCCTCTGTTTCGATAGCTTCGATGCTCTTGCAAATGGC
GAACCACATAGCACGTGATCACCCGATAATTGCCTAATGTTGTATCCTAGCAACACCGGCTTTTAGTGCG
ATGACTACGTGCACTTGTGGTTTCTGGCATTCATGACACTTTCAAAAATCGTACAAGGTGCACGCAGCTG
CGTTTGTCACGCATTCGGGCTTGTTACGACGTGTAATCGTAGCTATTTATCTACCGTACGTGAGCCTGTA
CCGGTGCACAGCATAACATAGCGTGTACAAAAGGCGCGCGTGTGGTCAGCTGATCTCTTGTACCGTTAAT
ACTCTGGCCTGCTCTGTTCTACATTGCCATTTGCGAACAGAAGCACAAGATGTATGTTTTGCAAGAGTTT
GCGTGAAAATAATCGAGGTGTGGTGGCTTGCAACGCTGAAAATACTATTTTTCCATCTCGACCAATCGGG
GACTCGAGGATCCGCGTGACGTAGTCCTTCCTCGATCGTTCGCCGCCTGCGATTGGTTGTTTCGAACGTG
ACCTCATGAATAGTAATTAGTTTGCTACGGCGTACATGTACTTTGAAGGAGGACCCCATATTGAGGTTGC
CGCGCTCCTCGGCGCCCGCTTTTCAGAGCTGCTTGTGTGTTCTACCGAGTCGGGTGCGTGCCATCCGCCT
GTGGTGCTGTAGTGCAAATGGAGAAGAGGATTGAATTGGAGAAGAGGGGTAAAAACCCCGAGCAGATACA
CGAGCTCAACCTCGACAACTGCAGGAGTACCGCCATCGTCGGCCTGACAGAGGAGTTCGTCAACTTGGAG
ACGCTCAGCCTCATCAATGTCGGCCTTACGAGCCTCAAGGGCTTCCCCAAGCTGCCCAACCTCAAGAAGT
TGGAGCTTAGCGACAACAGGATTTCTGGTGGCCTCAACCTGCTGCACGGGAGCCCCAAGCTCACACATTT
AAATTTGAGCGGGAACAAAATCAAGGGTCTGGAGACCTTGGATCCTCTGAAAGAGTTCAAGAACCTTAAG
AACCTTGATCTTTTCAACTGTGAAGTGACAAGCATTGAGAACTACAGAGACAGAGTCTTCGAACTTATCC
CGAGCTTGAAGTACCTGGACGGCTATGACCGAGATGAGAAGGAGGCAGAGGACTCCGAGGTGGACGACGA
AGATGGCAATGAGGAGGATGACGAGGAGAATGAAGTGGATGGGGAGGGAGAGAGTGATGAGGACGATGAA
GATGATGTAAACGTGGACGACGAAGATGATGAGGATGGGGAGGAAGATGACGACGTCGATGAGGAGGATG
GAGAAGATGGGGAGGAAGAGGTCGGTCTCGATTACCTTGAAAAAGACGACATTGATGAGGAAAGTGAAGG
AGATTTCTACAACCCATATGATGTGGATGATGACGAAGATGGAGATGAAAACGAGTCTCCAAAAGGCCAG
AAGCGAAAGCGGGAAGAGGAAGATGGGGAGGACTGAGGGAGGGTGGGAAGAGCACTCAACACAACTGGAA
CATCCCGGCATCTTTACTTTAGTTTTTTTTTATGTCTTCTTCATTGTCACCTGTAGACTTTGTACAGCGG
CAGAGAGAAACAAAATGCAACATCACAATCGCATCATCCTCGGTACCTGTCCCTAGCCTTTCTTCCCCCT
TGTTCATTTCATTTGTGAACAATTTTTTGTGTGTGTGTGTTTTAAAAAGCTTTTGCATGCATGGATTCCC
GTTAAGATGCCCACCCAGTTGCTTCGAGATTTGTATAGGTTGTTACTGCTAAACCAGATTTGATTTCTGG
CTACGACCGCAATCAATCGTGCTTACAGCCATGCACATTCCAAGAGCTGTGCTGTTACGGGCGCTTTTGT
CTTTTGCATCTGCGCCTTCCTCAATCTGAGGGACATCAAGTTAACTTTGAAAGCAGCGACACCAAAGTTG
GCCCAGAAAAGGGGGTGGCCCCTAGGGACAGTGCTGTGCAGGCACATCTACATCGTCATCCGCGTTTTTT
TTTTTCTTTCTCCTTGGCCCTCCCTTTTCTCGATTTGCTTTCAAAACATTGAATGAAATGCCTTTTATTG
TCGCATTGTTGGCTTCGAGGGAAGGGCCTGTGCGGGTGCACTGCACTTTTAATGTGAGCGACCGTGCGGG
CGCTTCCGTTGCGGCTGCCAGTCTTCTGTGGCACAGAGGTGCGCAGAAGTGTAAATTATCAGCAGGGGCT
TTGTCCAGTGTTTGGTTCAATTGTTAAATAAAGTTTTTACTGAAATGGCGAAAAAAAAAA

>lone star Aam ANP32A assembled mRNA short
AAATATCCCCTCGATGCTATGCCTGTTTTGCGGCAGAGCGTTTTCAGTAAGCGCGACATAGAGTTCGGCG
TGTCCTACGGTTCCGTTGCAACGGTGGCTGGCATGGCTCGCGCGGACTCCACACTGAATGCGACATTTTC
GAAGCAAGGTGTGCTCGGAAACGTGCCGAGGTGCCACGAAACGAAGACGAAAATTAGTTAGTGCTCGGCG
AACGAGGCAAGCCTGTAGTAACAAACAAACAGCAGCGATGGCGGCGTTTGGCGTTGCGGACTGTTTAGTC
GAGAACTCTCGGTATAAAGCAAGCGCTGATATCAACTAGAGAACATGCGAAAGGCACGGAGCTGCTTGTG
TGTTCTACCGAGTCGGGTGCGTGCCATCCGCCTGTGGTGCTGTAGTGCAAATGGAGAAGAGGATTGAATT
GGAGAAGAGGGGTAAAAACCCCGAGCAGATACACGAGCTCAACCTCGACAACTGCAGGAGTACCGCCATC
GTCGGCCTGACAGAGGAGTTCGTCAACTTGGAGACGCTCAGCCTCATCAATGTCGGCCTTACGAGCCTCA
AGGGCTTCCCCAAGCTGCCCAACCTCAAGAAGTTGGAGCTTAGCGACAACAGGATTTCTGGTGGCCTCAA
CCTGCTGCACGGGAGCCCCAAGCTCACACATTTAAATTTGAGCGGGAACAAAATCAAGGGTCTGGAGACC
TTGGATCCTCTGAAAGAGTTCAAGAACCTTAAGAACCTTGATCTTTTCAACTGTGAAGTGACAAGCATTG

AGAACTACAGAGACAGAGTCTTCGAACTTATCCCGAGCTTGAAGTACCTGGACGGCTATGACCGAGATGA
GAAGGAGGCAGAGGACTCCGAGGTGGACGACGAAGATGGCAATGAGGAGGATGACGAGGAGAATGAAGTG
GATGGGGAGGGAGAGAGTGATGAGGACGATGAAGATGATGTAAACGTGGACGACGAAGATGATGAGGATG
GGGAGGAAGATGACGACGTCGATGAGGAGGATGGAGAAGATGGGGAGGAAGAGGAGGAAAGTGAAGGAGA
TTTCTACAACCCATATGATGTGGATGATGACGAAGATGGAGATGAAAACGAGTCTCCAAAAGGCCAGAAG
CGAAAGCGGGAAGAGGAAGATGGGGAGGACTGAGGGAGGGTGGGAAGAGCACTCAACACAACTGGAACAT
CCCGGCATCTTTACTTTAGTTTTTTTTTTATGTCTTCTTCATTGTCACCTGTAGACTTTGTACAGCGGCAG
AGAGAAACAAAATGCAACATCACAATCGCATCATCCTCGGTACCTGTCCCTAGCCTTTCTTCCCCCTTGT
TCATTTCATTTGTGAACAATTTTTTGTGTGTGTGTGTTTTAAAAAGCTTTTGCATGCATGGATTCCCGTT
AAGATGCCCACCCAGTTGCTTCGAGATTTGTATAGGTTGTTACTGCTAAACCAGATTTGATTTCTGGCTA
CGACCGCAATCAATCGTGCTTACAGCCATGCACATTCCAAGAGCTGTGCTGTTACGGGCGCTTTTGTCTT
TTGCATCTGCGCCTTCCTCAATCTGAGGGACATCAAGTTAACTTTGAAAGCAGCGACACCAAAGTTGGCC
CAGAAAAGGGGGTGGCCCCTAGGGACAGTGCTGTGCAGGCACATCTACATCGTCATCCGCGTTTTTTTTT
TTCTTTCTCCTTGGCCCTCCCTTTTCTCGATTTGCTTTCAAAACATTGAATGAAATGCCTTTTATTGTCG
CATTGTTGGCTTCGAGGGAAGGGCCTGTGCGGGTGCACTGCACTTTTAATGTGAGCGACCGTGCGGGCGC
TTCCGTTGCGGCTGCCAGTCTTCTGTGGCACAGAGGTGCGCAGAAGTGTAAATTATCAGCAGGGGCTTTG
TCCAGTGTTTGGTTCAATTGTTAAATAAAGTTTTTACTGAAATGGCGAAAAAAAAAA
